# Aging Rhesus Macaque show tissue and sex-specific balance of drifting and coordinated miRNA programs

**DOI:** 10.64898/2026.04.27.720577

**Authors:** Shusruto Rishik, Erik Nutma, Nicole Ludwig, Ulrike Fischer, Tanja Tänzer, Stefanie Rheinheimer, Martin Hart, Laura Gröger, Johanna Meerjanssen, Matthias Flotho, Miguel A. Esteban, Jinte Middeldorp, Andreas Keller

**Author notes:** Correspondence should be addressed to Prof. Dr. Andreas Keller.

## Abstract

Macaque’s research centrality makes it critical to study their molecular aging. We accomplish this for their non-coding transcriptome by sequencing small RNA from 11 organs, with special focus on brain by including 24 brain regions, sampling males and females between ages 3-35 years. Heart, adrenal gland, corpus callosum and caudate putamen showed the most age-deregulated miRNA trajectories. The MIR-154 family, inside the imprinted, rejuvenation-associated *Dlk1-Dio3* cluster, was particularly vulnerable. Known age-associated miRNA families LET-7, MIR-29, MIR-17 and MIR-92 were strongly deregulated, with heavy dependence on tissue and sex. MiRNA genomic clusters deregulation was concordant within tissue-sex combinations, implicating upstream regulation rather than random noise. Cross-species comparison with mouse showed ancient miRNAs dominating age-deregulated trajectories. Deregulation direction in tissues-sex was conserved between species at family/cluster levels, but conservation substantially weakened at individual miRNA level. Thus, we mark a decisive step in translating miRNA aging trajectories between two heavily used model organisms.

**Key Findings:**

- Heart, adrenal gland, corpus callosum, caudate putamen are hotspots of miRNA age deregulation, with dramatic influence from sex.
- Non-brain organs show tissue specific miRNA change, with inconsistent overlap between tissues.
- Genomic clusters of miRNAs were found to be concordant in their age deregulation direction, dependent on tissue and sex, suggesting upstream regulation.
- The MIR-154 family, housed inside the heavily imprinted *Dlk1-Dio3* cluster and processed from the rejuvenation associated MEG3-MIRG host gene is prominently involved in both non-brain organs and brain regions.
- Concentration of age deregulation in evolutionarily ancient miRNAs across species implies regulatory program rather than epigenetic drift, involving MIR-154, LET-7, MIR-29, MIR-17 and MIR-92 families.
- Direction of change conserved between species at the family / genomic cluster level but diminished substantially at individual miRNA level.

## Introduction

Between the years 2015 and 2050, the global population over the age of 60 is expected to double to 2.1 billion while the population over 80 is expected to triple^1^. Improvement in health-span has not kept pace with increase in lifespan, resulting in many individuals suffering substantial age-related decline in quality of life. This shift places a significant burden on individuals and healthcare systems with people 85-and-above requiring up to 7 times more medical expenditure compared to individuals in their late-30s^2^. Therefore, there is growing interest in understanding aging as a biological process that can be intervened upon to increase human health span.

Mortality increases exponentially with age and can be modelled as an accumulation of multiple sources of damage over time^3^. However, finding these specific components and designing appropriate interventions represents an ongoing challenge. Major progress made so far has been summarized in the Hallmarks of Aging^4^ and includes somatic DNA damage accumulation with age^5^, loss of proteostasis^6^, deregulation of lncRNA^7^ and loss of CpG methylation^8^. Among the multiple layers of epigenetic regulation, miRNAs are included in the hallmarks of aging.

Since the identification of the first miRNA lin-4^9^, our knowledge of this class of non-coding RNA has exploded. MiRNA biogenesis, species conservation^10^ and 3’ untranslated region targeted downregulation of mRNA is well-established. In addition to fine-tuning target gene expression levels^11,12^, they cancel noise in randomly transcribed mRNA^13^ and canalize the transcriptome of cell types^14^. Deepening mechanistic understanding of miRNA targeting^15,16^ has opened up their application in iPSC reprogramming^17^, tissue-specific gene vector transcription^18^ and a recent breakthrough in the phase 1/2 treatment of Huntington’s Disease^19^. Atlas-level next-generation sequencing studies^20,21^ shows that the expression of miRNAs is highly organ-specific and serve as signatures of age-related diseases such as cancer^22^ and Parkinson’s disease progression^23^.

MiRNA expression change is widespread during aging, showing strong tissue-specific signals in human brains^24^, in mice full body organs^25^ and brain regions^26^. Multiple miRNA families such as MIR-29^25^, MIR-17^27^ and MIR-188^28^ have emerged as age-associated miRNA, alongside being involved in general stress response^29^. However, this information has not yet been translated to humans. While mice share miRNA targeting and expression patterns with humans, there are also large families of miRNA that are specific to primates^30^. Even though miRNA sequences themselves are often heavily conserved across mammalian genomes, miRNAs are often transcribed from poorly conserved long non-coding RNA host genes^31,32^ or vary in their genomic context such as transcription factor binding sites or their chromosomal position^33^. This difference is potentially contributing to the known cross-species translatability problem; models of Alzheimer’s/Parkinson’s disease can be cured in mice but findings do not translate to humans^34,35^. Given the different rates of primate and rodent aging, there are likely also differences in how miRNAs deregulate with age. Sampling otherwise healthy human organs and brain regions over the lifespan is not ethically possible. Thus, characterizing miRNA change with age for species closer to humans is important, in addition to understanding how murine findings translate between species. Macaques represent such a step between humans and mice, alongside being one of the most heavily used model organisms in medical research^36^.

While there are studies of miRNA expression in individual tissues for macaque^37,38^, aging is not localized and happens at a system wide level. As such, a comprehensive atlas of miRNA expression from organs and brain regions for monkeys of different ages is lacking. Therefore, in this study we perform bulk small RNA sequencing of multiple major body organs and brain subregions for both male and female monkeys across the lifespan for rhesus macaques. Our study represents the largest such cohort and gives a peek into tissue and brain-region specific deregulation of non-coding RNA with age. We also leverage our previous expertise from the non-coding arm of Tabula Muris Senis to explore the cross-species translatability of age deregulation.

## Results

We sampled 13 organs in total, including brain. Since we previously found brain regions to age differently in our mouse aging study^26^, we additionally sectioned the brain. To avoid confusion, we refer to the samples as brain region and non-brain organ. We have 283 samples from 28 individual macaques from two closely related species (23 from *Macacca mulatta* and 5 from *Macacca fascicularis*) across 11 non-brain organs (stomach, spleen, colon, lung, kidney, heart, limb muscle, adrenal gland, thyroid, liver and gallbladder) [Figure 1a]. Both male (age range 5.35-22.53 years) and female monkeys (age range 8.31-34.84 years) represented [Figure 1b], to explore the impact of sex in aging. We note the large difference in ranges stemming from two female monkeys of advanced age. The inclusion of two species is motivated by the knowledge that miRNA expression patterns can be different even in *M. musculus* strains^39,40^. We wanted to see if strain-level differences in *M. musculus* had a parallel in different species of macaques.

**Figure 1:**
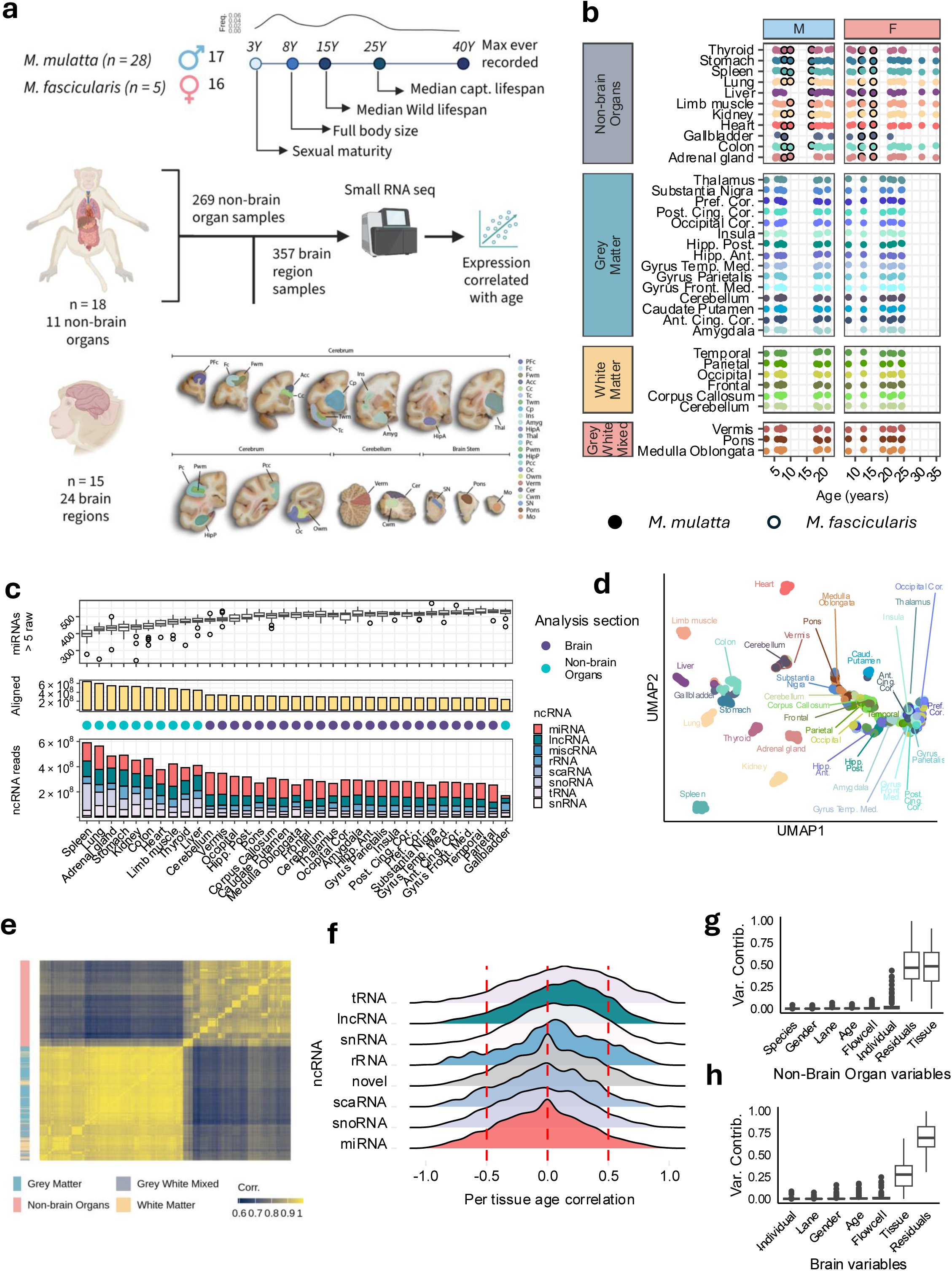
Data distribution a) Study setup and macaque species, sex, tissues, brain regions and age along with typical lifespan of macaque monkeys. Macaque anatomy and NGS illustrations created using BioRender. Note, anatomy cartoons from Biorender are for illustration purposes only and does not represent actual Macaque anatomy. Only brain regions marked with color correspond to photographs of actual samples. b) Sampling distribution of macaque monkeys over sex, age, organs and brain regions. Each timepoint corresponds to a sample from an individual. Bordered dot corresponds to *M. fascicularis* while un-bordered dot corresponds to *M. mulatta*. c) Distribution of miRNAs passing the detection threshold, number of reads aligned to macaque genome, and the fraction of each non-coding RNA class included in the aligned reads. d) UMAP embedding of organs and brain regions based on miRNA rpmm expression passing filtering threshold. e) Heatmap of pairwise correlation of each sample, based on rpmm normalized expression values of miRNA passing the filtering threshold. f) Distribution of spearman rank correlation for each class of non-coding RNA, based on rpmm normalized expression. Only non-coding RNA passing filtering threshold were kept. g) Variance contribution of variables Species, Gender, Age, Tissue, Lane, Flowcell, Individual and Residuals on miRNA passing detection filter for organ subset of data based on a linear mixed model. h) Variance contribution of variables Individual, Lane, Gender, Age, Flowcell, Tissue (Brain Region) and Residuals on miRNA passing detection filter for brain samples based on a linear mixed model.

For the brain regions, we have 357 samples from 15 individual monkeys from the species *Macacca mulatta* from 8 males (age range: 2.5 - 21.42) and 7 females (age range: 7.92 - 24.42). We sampled 24 brain regions (Grey matter: occipital cortex, gyrus temporalis medius, gyrus frontalis medius, anterior cingulate cortex, gyrus parietalis, insula, prefrontal cortex, cerebellum GM, posterior cingulate cortex. White matter: occipital WM, cerebellum WM, corpus callosum, frontal WM, parietal WM, temporal WM along with vermis, amygdala, hippocampus anterior, medulla oblongata, hippocampus posterior, thalamus, caudate putamen, pons and substantia nigra) [Figure 1a, b]. We know that certain parts of the brain are more vulnerable to age-related degeneration, marked by deregulation of mRNA and miRNA. Therefore, we emphasize looking at brain regions to prioritize those especially deregulated with age.

### Overview of the data

In total we sequenced 17 billion small RNA reads. Figure 1c shows the total number of aligned reads to the macaque genome distributed between 60 million and 20 million among non-brain organs/brain-regions. Per sample, the number of reads aligned had a median of 20 million (interquartile range of 4.74 million) [Supplementary Table 1]. Of note, the brain region samples had lower aligned reads in comparison to non-brain organs, except for gallbladder [Figure 1c]. However, the number of miRNAs reads detected in brain regions at > 5 raw count per sample is consistently larger than non-brain organs. At a per sample level, we have a median of 496 miRNAs detected (interquartile range of 56). After filtering stably expressed miRNA (see Methods for threshold), we were left with 665 miRNAs out of 990 miRNAs annotated in miRbase.

Interestingly, median expression *of M. mulatta* and *M. fascicularis* were highly similar between non-brain organs in both females and males [Supplementary Figure 1a, 1b]. This is despite *M. mulatta* having a somewhat longer lifespan compared to *M. fascicularis* (median age at death: 6.93-9.84 years for *M. mulatta* vs. 7.89-10.49 years)^41^. Given that the age range of *M. fascicularis* individuals are small compared to *M. mulatta*, we opted to include them in the aging trajectory analysis and refer to the full set as macaque.

We perform a UMAP embedding of the samples using miRNA expression and color by tissue and see that non-brain organs and brain-regions cluster separately [Figure 1d]. colon, gallbladder and stomach form a central cluster while the brain-regions cluster away from the non-brain organs, being split by the type of brain matter. Plotting a heatmap of the pairwise correlation highlights this in Figure 1e. Interestingly, the non-brain organs and brain regions themselves show different within-cluster similarity; the within-group similarity for brain-regions is greater than for non-brain organs.

Since the impact of age is overwhelmed by the tissue, we calculate spearman rank correlation coefficient and show the distribution of the different classes of non-coding RNA in Figure 1f. Sorting by median correlation coefficient, we see tRNA, lncRNA and snRNA are skewed positively. Meanwhile miRNA is skewed negatively. Fitting a linear mixed model using the biological and technical variables we see the variance contribution of each variable to the miRNA in Figure 1g and 1h. As expected, for non-brain organs,

“Tissue” is contributing most of the variance. Meanwhile, for non-brain regions, “Residuals” are the biggest contributor followed by “Tissue”. Interestingly, in the non-brain organs, the macaque “Individual” variable is contributing the third most variance, while being in the last position for brain-regions. The presence of “Flowcell” appearing before “Age” led us to investigating if this contributes to a major batch effect. Plotting this on a UMAP in Supplementary Figure 1c, we see this overlapping with the non-brain organs and brain subsections. Each tissue forms a relatively tight cluster without obvious influence from “Flowcell”. Similarly, obvious separation based on “Gender”, “Lane” and “Species” could not be detected from UMAP embeddings [Supplementary Figure 1d, 1e, 1f]. Of note, gallbladder appears to have a low number of reads aligning to miRNA while simultaneously having one of the highest number of miRNAs detected. Therefore, large signals originating from gallbladder were deprioritized.

### Age deregulated miRNAs show systemic changes linked by known circulating miRNAs in plasma and serum

To identify similarity in miRNA age deregulation across the organism, we investigate the tissue/sex combinations. MiRNAs are classed into families based on sequence and seed region similarity, with the latter determining their targeting repertoire^10^. Relying on this, we ranked the total number of trajectories from each family in Figure 2a. We can see a skew towards negative deregulation for all miRNA families for all tissue-sex combinations with 4,102 and 2,529 trajectories in the negative and positive directions [Supplementary Table 2]. Each miRNA deregulated in a tissue in a specific gender is counted as a trajectory. Additionally, we color miRNAs based on known presence in serum or plasma in macaques.

**Figure 2:**
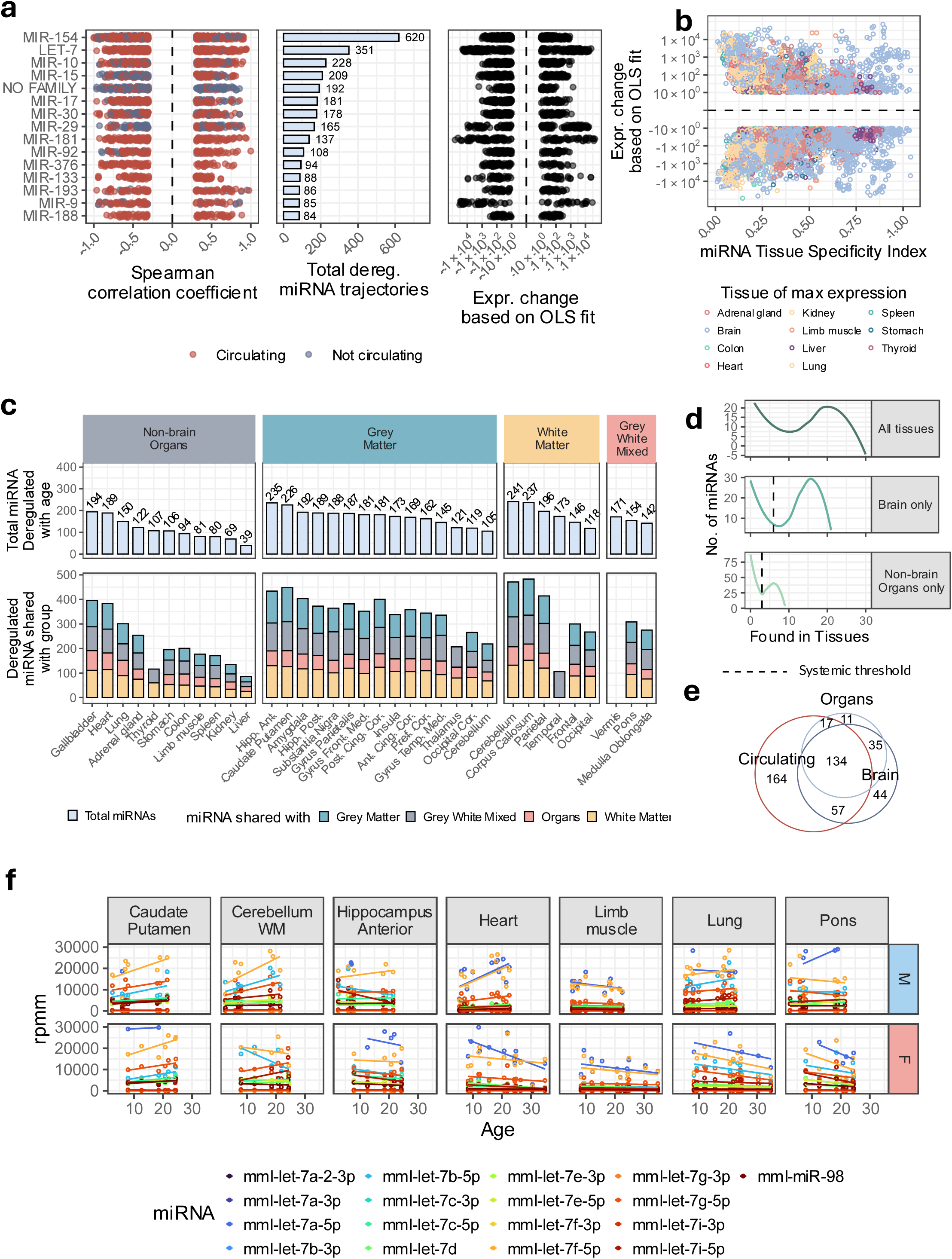
Systemic changes of miRNA with age in organs and brain regions a) Distribution of spearman rank correlation coefficient for members of the miRNA families that are most frequently occurring in the datasets, along with their count, and the magnitude of expression change over the lifespan. b) Scatterplot of tissue specificity index of miRNA with the magnitude of expression change over the lifespan. Brain regions have been collapsed into a single organ, to prevent deflation of TSI score. The miRNAs are colored based on the tissue in which the expression is maximum. c) Total number of miRNAs passing the age-related deregulation effect size threshold split by non-brain organs and brain organs. Shared miRNA counts show number of miRNA each organ/ brain region is sharing with at least one other organ/ brain region from the categories Organ, Grey Matter, Grey White Mixed and White Matter. d) Distribution of number of tissues and brain regions of miRNAs passing age-related deregulation effect size thresholds. e) Overlap of miRNAs deregulated with age from organs and brain regions with miRNAs detected in circulating fractions (plasma or serum fractions) f) Age trajectory plot of LET-7 miRNA family; the family with largest overall count of miRNAs deregulated with change and the greatest range of deregulation magnitude.

MiRNA families with most deregulated trajectories are MIR-154, LET-7, MIR-10 and MIR-15 [Figure 2a]. Interestingly, singular miRNAs are included in the top 5 families with the most trajectories. Notably, even though the distribution of spearman rank correlation distribution is similar across the families, the absolute change over time differs. While MIR-154 (Belonging to one of the largest genomic clusters of miRNA in macaque, mouse and humans, *Dlk1-Dio3* cluster) have almost twice as many trajectories as LET-7, LET-7 has a much broader distribution of change over age. The larger changes are prioritized in this manuscript as they are less likely to be technical artefacts [Supplementary Figure 1g]. Interestingly, except for MIR-15, MIR-29 and the miRNAs without an assigned family, a majority (101 of 140) miRNA members of the top families are also found in the circulating set [Figure 2a, Supplementary Figure 1h].

Since miRNA are highly tissue specific^21^, we wanted to see if tissue specificity was related to age related deregulation[Figure 2b]. Interestingly, miRNA aging trajectories are enriched for miRNAs with TSI values less than 0.75 (91.8%, 6089 trajectories out of 6631 for miRNAs with TSI < 0.75) [Supplementary Table 3]. Previous reports show evolutionarily established miRNA become more broadly and highly expressed^30^. With increasing miRNA tissue specificity, there is a decreasing trend in absolute magnitude of change. Deregulation trajectories involving miRNAs that are most heavily expressed in non-brain organs dominate the range TSI < 0.5 (non-brain representing 3143 out of 4666 trajectories). Meanwhile for TSI > 0.5, the miRNA involved are most highly expressed in the brain (brain representing 1201 out of 1965). This suggests a partitioning of the miRNAs involved in age deregulation between brain and non-brain organs and warrants a separation of the analysis for these miRNA trajectories.

Interestingly, we found miRNAs deregulated in virtually all the tissues we examined [Figure 2c]. Given the poor sequencing depth of gallbladder, we bypass it and state that heart, lung and adrenal glands as the non-brain organs most deregulated with 189, 150 and 122 miRNAs respectively. In the brain region, White matter appears to be more deregulated in comparison to grey matter with cerebellum and corpus callosum having 241 and 237 miRNAs. Meanwhile hippocampus anterior, caudate putamen and amygdala are the grey matter brain regions with the greatest deregulation with 235, 226 and 192 miRNAs. Interestingly, mixed grey-white matter from the brain stem have comparatively lower number of deregulated miRNAs at 154 and 142 miRNAs from pons and medulla oblongata.

Given that only 665 miRNAs are considered across the whole dataset, there must be overlapping miRNA deregulation. We tabulated the number of miRNAs that were shared with at least one member of the non-brain organs, Grey Matter, White Matter and Grey White Mixed matter groups for each tissue to identify if certain groups were sharing more heavily with others. However, from Figure 2c, we see that the number of miRNAs shared is proportional to the number of deregulated miRNA. The only exceptions are thyroid, which does not share miRNAs with Grey White Mixed and Grey Matter, temporal lobe of the white matter which only shares miRNAs with non-brain organs, thalamus grey matter which does not share anything else with any other grey matter, and vermis, which doesn’t share miRNA with any other tissue at all.

Given the large number of miRNAs shared between each tissue and at least one other from each category, we wanted to quantify the number of miRNAs that are shared between all tissues, within brain regions and within non-brain regions. Looking at the number of miRNAs shared between tissue in Figure 2d, we see bimodal distributions, with a set of miRNAs that are specific and another that are shared. When we consider all tissues, we have ∼20 miRNAs for both specific and systemic. Meanwhile, for brains this number is larger with ∼30 miRNAs for brain-region specific and brain-region shared miRNAs respectively. For non-brain organs, the tissue specific miRNA count is much larger, at ∼80 miRNAs, while shared miRNAs are about 40. Interestingly, as the number of non-brain organs / brain-regions increases, the shared miRNAs peak and then collapse to zero. While we do not have blood in our aging dataset, we see that a majority of (251 out of 374 unique miRNAs) that are deregulated have been previously reported in at least one of three circulating miRNA datasets [Figure 2e].

Using LET-7, the miRNA family that is the most widely deregulated (351 total trajectories) with the largest range of absolute expression change (between-20,727 rpmm and 23917 rpmm) to highlight how they change over time in Figure 2f. We note that directionality is shared neither between male and female nor between different tissues. Thus, the miRNA deregulation landscape seems to be heavily context dependent, warranting a deeper look.

### Sex-dependent miRNA clusters deregulated with age in non-brain organs

Looking at the frequency of the deregulated miRNA counts for each non-brain organ in each sex we see that there is a clear difference based on sex, not only in terms of the number of miRNAs deregulated in each tissue, but also the directionality of change. Heart is the tissue with the highest number of deregulated in both male and female, with 122 and 155 miRNAs respectively [Figure 3a]. Given that there are 189 miRNAs in total that are deregulated, there is some overlap guaranteed, however, there are no miRNAs changing in the same direction in both genders [Supplementary Figure 1i]. Similarly, lung is heavily deregulated in female monkeys with 132 miRNAs, but not as severely deregulated in males, with only 40 miRNAs. Looking at the overlaps between male and female, we see that the downregulated miRNAs are shared between sexes, while the upregulated miRNAs are male-specific [Supplementary Figure 1i].

**Figure 3:**
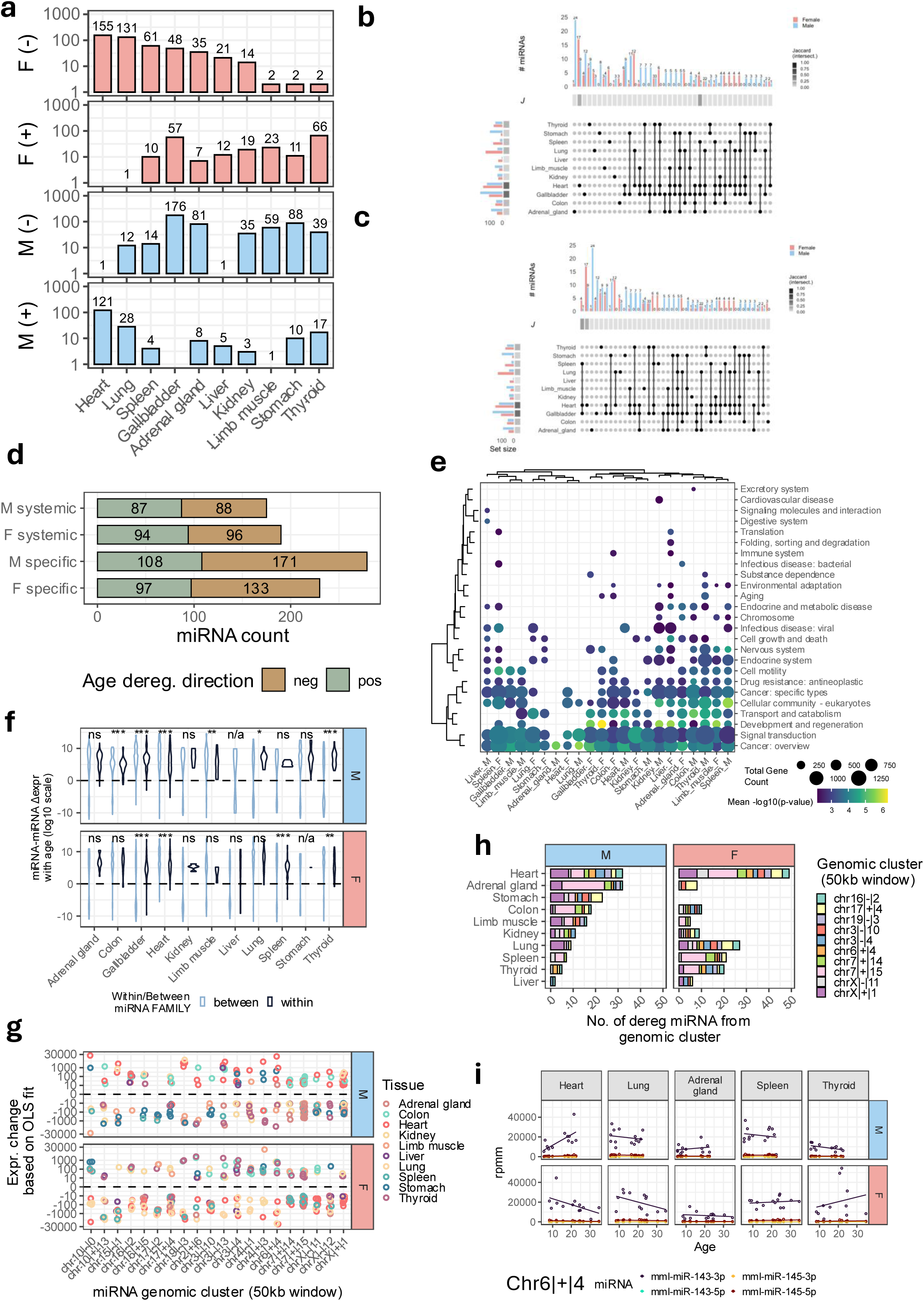
Sex-dependent systemic and specific changes in macaque organ miRNAs. a) Number of miRNA deregulated with age for females (F) and males (M) in the positive (+) and negative (-) directions. b) Overlap of miRNAs deregulated with age within organs for female and male monkeys, sorted by frequency in set overlap. c) Overlap of miRNAs deregulated with age within organs for female and male monkeys, sorted by jaccard index of overlap for each set d) Number of miRNAs that were found to be shared across organs (systemic) and those that were tissue specific (specific) in male (M) and female (F). Color represents if the change is in the positive versus negative direction. e) Pathway subcategories overrepresented by targets of age-deregulated miRNAs for organ-tissue combination. f) Concordance distance distribution of miRNA-miRNA pairs within the same genomic cluster versus between different genomic clusters. ns corresponds to non-significant. n/a corresponds to not applicable due to lack of miRNA in one of the groups. *, **, *** corresponds to p-value less than 0.05, 0.01, 0.001 respectively. g) Magnitude of deregulation over lifespan for miRNAs from each non-brain organ, split according to genomic cluster. Top 20 miRNAs were selected based on mean absolute change of miRNAs in that cluster. h) Number of miRNAs from each genomic cluster deregulated in each organ. Only the top 10 most frequently occurring genomic clusters are shown. i) Age trajectory of members of a selected cluster (chr6|+|4).

Given the dramatic between-sex differences, we wanted to check in more detail the within-sex overlaps for the non-brain organs [Supplementary Table 4, 5]. Our results from Figure 3b align with the previous bimodal distribution in Figure 2d; sorting the set overlaps by number of miRNAs prioritize single-tissue sets, suggesting a stronger tissue-specific signal. Additionally, the overlaps are typically not shared between the sexes; looking at Figure 3c which sorts the sets by Jaccard index of overlap, we see a skew towards tissue specific miRNAs, with the Jaccard index value falling off from 0.25 after the first two overlapping sets. If we look at the unique number of miRNAs that are systemic versus tissue-specific in male and female, [Figure 3d]. Thus, while the set of miRNAs are limited, their deregulation appears to be context specific.

To check if the miRNA deregulated in each non-brain organ within each sex are modulating different pathways, we assigned ontology using predicted targets of the deregulated miRNAs [Supplementary Table 6]. At the pathway level, we see both tissue specific clusters deregulated and pathways that are shared across tissues [Supplementary Figure 1j]. Commonly known pathways associated with aging such as the Ras signalling pathway(mcc04014), AGE-RAGE signalling pathway (mcc04933), mTOR signalling pathway (mcc04150), Wnt signalling pathway (mcc04310) are represented in a non-tissue-specific manner. When we take the pathways collapsed by subcategory, sets overrepresented by the targets of these miRNAs converge onto cancer (22/ 22 tissue-sex combinations), signal transduction (21 / 22 tissue-sex combinations), development and regeneration pathways (15 out of 22 tissue-sex combinations) [Figure 3e]. No obvious clustering is noticeable between the non-brain organs or sexes. Unfortunately, at the present state of the field, narrowing down to experimentally validated targets is not possible due to the dearth of experimentally validated targets for macaque miRNAs.

Since miRNAs are often genomically clustered with the synteny conserved among species, we suspect deregulation among these clusters would represent coordinated programs changing with age. Meanwhile, non-clustered miRNAs being deregulated are unlikely to constitute a strong regulatory signal, representing the noisy part of miRNA deregulation with age. Therefore, we test this hypothesis by seeing if deregulation within-clusters is concordant compared to between-clusters [Figure 3f, Supplementary Table 7]. We split it up by sex and non-brain organ as we suspect these two strongly contextualize upstream regulation. Interestingly, the between-cluster distances tend to be centered around 0, while the within-cluster deregulation appears to be centered in positive direction, e.g. male heart has a median of 7.94 (IQR 6.00) and female heart has a median of 7.41 (IQR 4.85). The distributions for certain tissues were found to be statistically different between within-cluster and between-cluster, e.g. colon has a p-value <0.001 in male but not in female. Since the distance was defined such that miRNA-miRNA pairs having matching directions would have positive distances, it suggests that the genomic clusters are deregulated towards shared directions, further suggesting coordinated upstream regulation of the clusters. However, since some tissues such as liver and stomach show no clustered deregulation in male, female respectively, it highlights that not all tissues are undergoing coordinated expression change for miRNAs.

Zooming into the concordant clusters with the largest changes in expression over the lifespan, we see that the concordance holds within tissue and with gender [Figure 3g]. The within-cluster changes vary, with certain clusters like chrX|+|1 showing a change between-1000 and 100 absolute rpmm change, while others like chr6|+|4 have a larger range spanning between-30,000 and 30,000 rpmm. While concordance holds within the context, Figure 3g suggests that different clusters might be deregulated in a tissue/gender specific way. Therefore, we count the top genomic clusters with deregulated members in each tissue/gender in Figure 3h. We see that heart is the only tissue with both genders showing coordinated changes (Total trajectories in Heart is 277 with 122 coming from males and 155 coming from female, with 79 and 87 coming from clusters with >= 2 miRNAs deregulated). However, the clustered deregulation in other non-brain organs is almost complementary between male and female; adrenal gland (43 out of 89), stomach (50 out of 98), colon (20 out of 50), limb muscle (21 out of 60) being deregulated in males, while lung (71 out of 132), spleen (35 out of 71) and thyroid (39 out of 68) being deregulated in females [Figure 3f]. Two of the clusters with the largest deregulated miRNAs chr7|+|15 which correspond to the *Dlk-Dio3* region housing the MIR-154 cluster, along with chrX|+|1 which is housed inside the *CLCN5* gene. It should be noted that while the MIR-154 contains 34 miRNA precursors, and the direction of expression tends to be concordant, it is not the case that every member generates mature miRNAs (e.g. in female heart, 10 out of 34 precursors have generated a mature miRNA, 17 mature miRNAs have been generated in male adrenal gland) suggesting a potential involvement of downstream processing change.

We chose Chr6|+|4 to illustrate the change of miRNAs inside a cluster over age. The cluster Chr6|+|4 has miRNA precursors mml-mir-143 and mml-mir-145 near a novel lncRNA in macaque (ENSMMUT00000071187), while corresponding human homologues are inside a dedicated host lncRNA (CARMN). Deregulation in representative non-brain organs depend upon the tissue and the sex. While expressions of the cluster members are concordant, they show a large difference in the mature miRNA expression levels derived from each precursor, e.g. mml-miR-143-3p has increased 18,427 rpmm in male heart, but mml-miR-143-5p change is below the threshold of detection. Therefore, even though both arms are derived from a single precursor, one arm has changed expression while the other has not changed detectably. This additionally hints at differences in downstream processing of mature miRNA, which might be altered with age.

### Brain shows diffused aging with MIR-154 cluster as particularly vulnerable

In the brain, White matter stands out as having the greatest number of deregulated miRNAs, with corpus callosum showing 198 miRNAs downregulated in females and 140 downregulated in males, second only to Hippocampus anterior (154) [Figure 4a]. The next most deregulated regions in females are the white matter in cerebellum (155), caudate putamen (154) and vermis in grey-white mixed matter (147). For males, the pons (95) and prefrontal cortex (87) are notably deregulated. If we look at the miRNAs that are deregulated in the same direction in males and females, we see a greater proportion of sharing in brain regions compared to non-brain organs. Specifically, for corpus callosum, there is the largest number of shared miRNA (106 miRNAs in the negative direction) [Supplementary Figure 1i]. Looking at it on a per tissue basis, females have miRNA that are primarily in the negative direction (median no. of trajectories for males and females are 30 and 50), while males have miRNA primarily in the positive direction (median no. of trajectories for males and females are 38 and 12).

**Figure 4:**
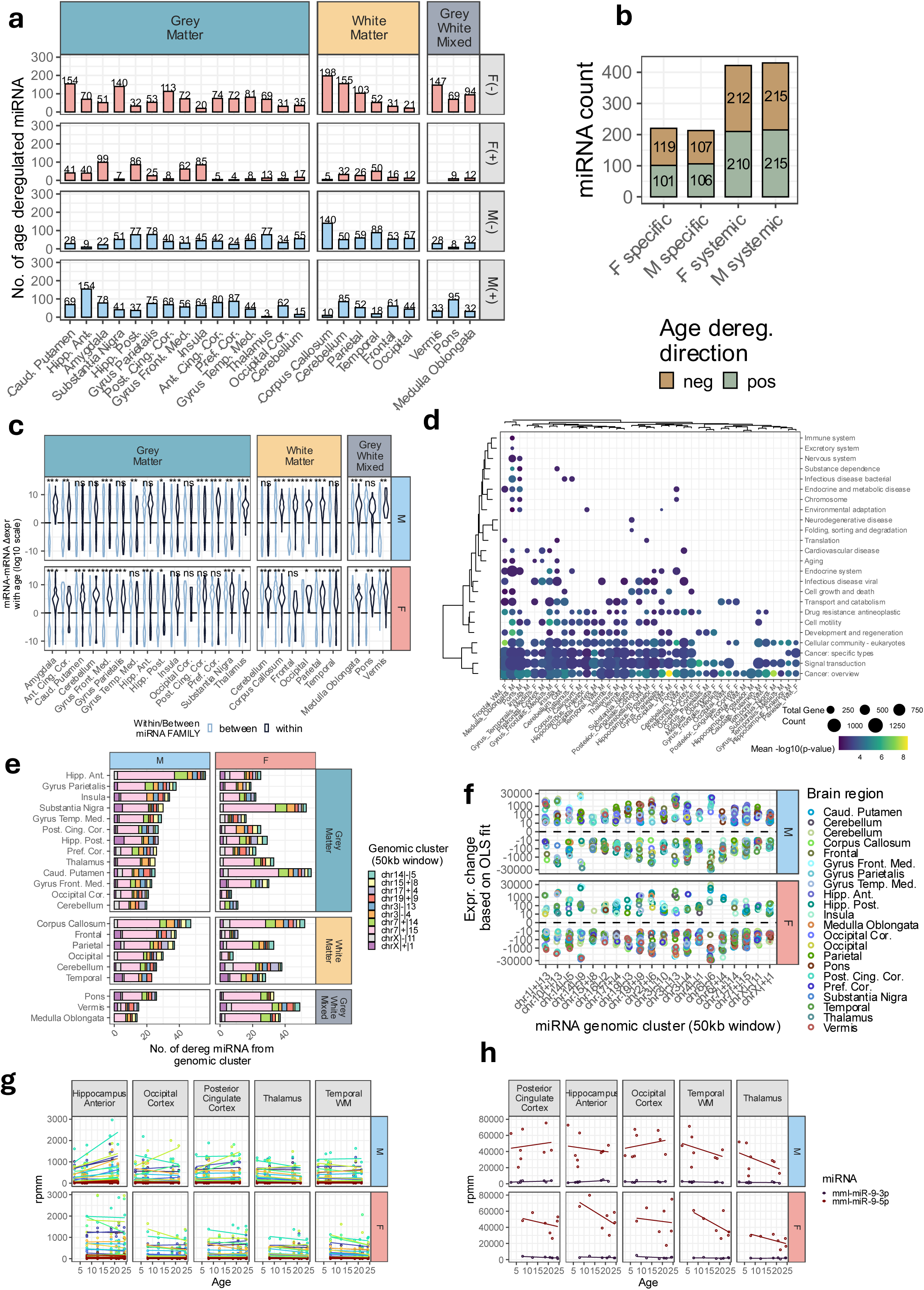
Sex-dependent brain region specific and diffuse miRNA age deregulation. a) Number of age-deregulated miRNA for each brain region, grouped by grey matter, white matter and grey-white mixed. Also split by male (M) and female (F), with deregulation going in positive (+) / negative (-) direction. b) Number of age deregulated miRNAs that are brain region specific versus found deregulated in a diffuse manner across brain region. c) Concordance distribution of miRNA change and direction within genomic cluster and between genomic clusters for each brain region, split by type of brain matter. ns corresponds to non-significant. n/a corresponds to not applicable due to lack of miRNA in one of the groups. *, **, *** corresponds to pvalue less than 0.05, 0.01, 0.001 respectively. d) Pathway subcategory overrepresentation using mRNA targets of age deregulated miRNAs in each brain region-gender combination. e) Number of age deregulated miRNAs in top 10 most frequent genomic clusters grouped by type of brain matter. f) Magnitude of age deregulated miRNAs for brain regions, split by genomic cluster. Only top 10 genomic clusters ordered by mean absolute change have been plotted. g) Genomic clusters that are most frequently deregulated with age (chr7|+|14 and chr7|+|15 corresponding to MIR-154) across 5 representative brain regions, split by male and female.h) Aging trajectory of nervous system specific miRNA miR-9 across 5 brain regions with largest deregulation.

If we look at miRNAs that are brain-region specific and systemically deregulated across the brain, there are approximately twice as many systemic deregulation as region-specific deregulation [Figure 4b]. Interestingly, here the directional skew between genders collapse. To determine if both male and female are having clustered expression in the brain regions, we plotted the concordance of clusters for the deregulated miRNAs as well [Figure 4c]. The within-cluster concordance is not as clear for brain regions as for non-brain organs. For example, while Corpus callosum is concordant in male and female. White matter cerebellum is concordant in females but highly discordant in males, despite both having similar number of deregulated miRNAs (187 in females and 125 in males). To see if the deregulated miRNAs are converging on identical pathways, we look at the pathways associated with the targets of the deregulated miRNAs. At the pathway level, we see certain inflammatory pathways such as T cell receptor signalling pathway (mcc04660), IgSF CAM signaling pathway (mcc04517) and pathways seen in non-brain organs such as Ras Signaling pathway (mcc04014), TGF-beta signaling pathway (mcc04350) overrepresented [Supplementary Figure 2a]. Again, grouping by subcategory results in cancer and signal transduction showing up without a specific focus on any brain region (40 out of 48 and 37 out of 48 sex-brain region combinations respectively) [Figure 4d].

The range of miRNAs generated from the top 15 genomic clusters (between 0 and ∼50 on Figure 4e) is small in comparison to the number of deregulated miRNAs. When considering miRNAs in clusters with at least 2 or more members, they become more comparable to non-brain organs. E.g. hippocampus Anterior in males have 163 miRNAs deregulated in total, with 89 in such clusters. Within these clusters, again it seems that chr7|+|15 is dominating the stacked bar plots across almost all brain regions with median number of members 9 (IQR =13) and 13 (IQR = 8) for females and males respectively [Figure 4e]. Plotting the expression change in the clusters and coloring by brain region highlights the concordance of the clusters within the regions [Figure 4f].

This cluster corresponds largely to the MIR-154 cluster, and to highlight the change we plot the trajectories of the members in Figure 4g. The plot highlights what we saw in Figure 2a: while the change over the life span is modest, the sheer number of miRNAs involved in this cluster (310 and 329 trajectories for females and males respectively) along with the clusters generally high expression makes it a locus of importance. However, as with non-brain organs, all members of this cluster do not get deregulated in all tissues.

To contrast the change of a set of miRNAs from the same genomic cluster, we looked at a central nervous system-specific miRNA precursor that is located on 3 different chromosomes. Mml-miR-9-5p shows identical direction change for hippocampus anterior, temporal WM and thalamus but differing direction in posterior cingulate cortex and occipital cortex for the sexes. While the expression of the 5’ end does change, the ratio of the 5’ to the 3’ mature miRNA does not, suggesting here that arm selection is not something dramatically impacted by aging.

### Cross-species age deregulation conserved at family level but diverging at individual miRNA level

Estimating cross-species translatability remains an open problem: while mice make a convenient model organism to perform experiments, specific results are often not translatable to humans. For example, while both mice and humans have miRNAs and RISC complexes to degrade mRNA, a majority of miRNA are not homologous in sequence between the two organisms. Perturbing a miRNA to obtain a phenotypic outcome in mice will not necessarily obtain the same outcome in humans. As such, we wanted to quantify the degree to which the findings of prior miRNA age trajectories from the Tabula Muris Senis project agree with macaque monkeys. It also serves as a validation to check if the concordant clustered deregulation patterns / tissue and sex-specific deregulation pattern seen in macaque also translate. Since sampling density for the mouse dataset is 3 times larger than the macaque data, it will also highlight what is potentially a sampling error [Supplementary Figure 2b].

Based on the annotations in miRbase and mirgenedb, we see only about half of the miRNAs in the miRbase annotations for *M. mulatta* are homologous to *M. musculus,* meanwhile only one-fourth of *M. musculus* miRNAs have homologues [Figure 5a]. Since miRNA tends to be tissue-specific, we observe the tissue specificity index distribution for *M. mulatta* and *M. musculus* in Figure 5b [Supplementary Table 8, 9]. The conserved and un-conserved miRNAs show two separate tissue-specificity index profiles with older, conserved miRNAs showing broad expression while un-conserved miRNAs show more tissue specific expression. If we consider the miRNAs that are conserved, the tissue specificity index has a strong correlation (spearman rho = 0.854, pvalue = 1.68×10^-139^) [Figure 5c]; the higher the tissue specificity index, the higher the probability of matching tissue of maximum expression. To identify which tissues are showing this kind of matching maximum expression, we additionally color this plot by Tissue of max expression in Supplementary Figure 2c. Brain, adrenal gland, heart and limb muscle are highly represented in the miRNAs that have matching TSI and tissue of maximum expression (39, 5, 5 and 4 miRNAs respectively out of 59 miRNAs with a TSI above 0.75 with matching tissue of max expression).

**Figure 5:**
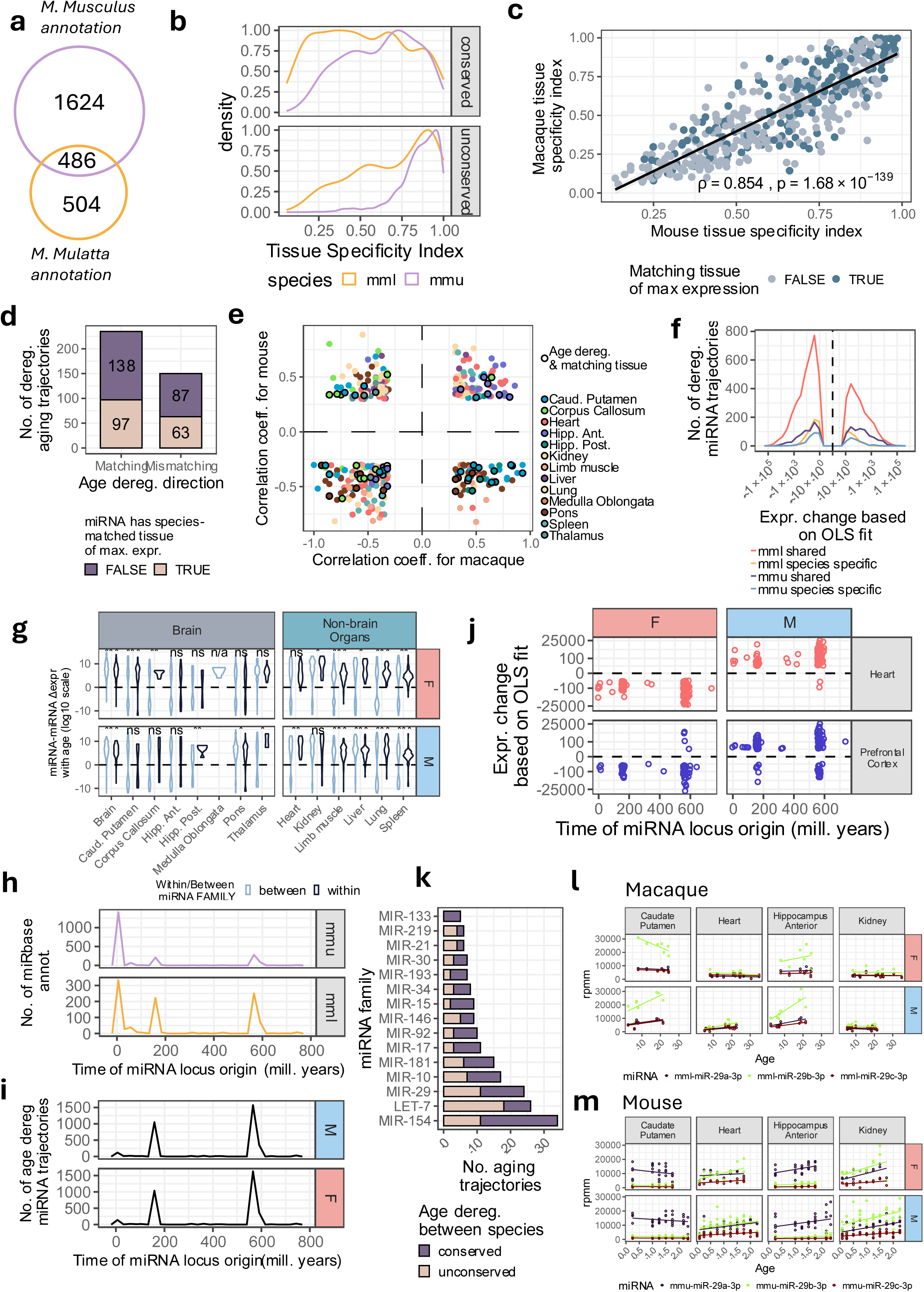
Correspondance of miRNA aging trajectory between mice and macaque. a) Overlap of homologous miRNAs between *M. musculus* and *M. mulatta*. b) Distribution of tissue specificity indices for conserved and unconserved miRNAs from 5a, colored by *M. mulatta* (mml) and *M. musculus* (mmu). c) Scatterplot of relationship between *M. mulatta* and *M. musculus* tissue specificity indices for miRNA that are homologous from 5a. If the tissue of maximum expression matched between species, the miRNA is colored as per legend. d) Frequency of miRNA that are matching in terms of deregulation direction (note, tissue of deregulation is not considered here) across lifespan and split by if miRNA has matching tissue of maximum expression based on 5c. e) Correlation coefficient scatterplot of miRNAs deregulation with age in *M. musculus* and *M. mulatta*. f) Distribution of expression change based on OLS fit for *M. musculus* (mmu) and *M. mulatta* (mml) miRNA that are shared across species and those that are species-specific. g) Concordance distance distribution for *M. musculus* miRNA brain and non-brain organs harmonized with *M. mulatta* brain and non-brain organs. ns corresponds to non-significant. n/a corresponds to not applicable due to lack of miRNA in one of the groups. *, **, *** corresponds to pvalue less than 0.05, 0.01, 0.001 respectively. j) Expression change for *M. mulatta* Heart and Prefrontal Cortex for miRNA loci and their corresponding time of origin (millions of years). h) Distribution of miRbase annotations for *M. musculus* (mmu) and *M. mulatta* (mml) over the time of miRNA loci origin (millions of years). i) Distribution of deregulated miRNAs for *M. mulatta* over time of miRNA loci origin (millions of years). k) Top 15 miRNA families from homologous miRNAs that have the greatest number of age deregulated trajectories for *M. mulatta*. Color differentiates miRNAs that have matching direction of deregulation between species. l) Line plots highlight miRNA change for members of MIR-29 family for representative brain and non-brain organs for *M. mulatta*. m) Line plots highlighting miRNA change for members of MIR-29 family in *M. musculus* for matching brain, non-brain organs.

Since there are matching tissue-specific expression patterns between macaques and mice, we wanted to explore if these miRNAs are also showing a conserved direction of deregulation with age in Figure 5d. Interestingly, there are more miRNA trajectory that have matching direction with age than mismatching (235 vs. 150). However, within each category, there are more miRNAs that are mismatching in terms of their tissue of max expression. Figure 5e highlights the divergence in more detail, with hippocampus anterior, corpus callosum and pons being among the tissues with trajectory directions also matching alongside tissue of maximum expression. While correlation coefficient calculates the strength and direction of association, it does not calculate how large the absolute change is over the course of the lifespan. We see peaks for species-specific and shared trajectories from both mouse and macaque between 10^1^ and 10^2^. There is a noticeable skew toward the negative with miRNAs that are shared between species showing larger changes, e.g. species-shared trajectories in macaque have a negative peak of around ∼800 trajectories compared to a peak at ∼400 trajectories in the positive direction [Figure 5f].

As we found concordance within clustered miRNA expression in macaque monkey, we wanted to see if this pattern also held in mice [Supplementary Table 10]. Interestingly, the pattern holds, with non-brain organs showing strong concordance with matching directions (5/6 showing significantly different distributions compared to between family groups). Brain, on the other hand shows a more diffuse deregulation pattern with miRNA pairwise distance both matching and mismatching (3/8 and 5/8 significantly different distributions for female and male respectively) [Figure 5g]. In terms of sex-differences, corpus callosum, hippocampus posterior, thalamus, kidney, and limb muscle show some of the most obvious difference in terms of within/ between miRNA cluster distribution, again suggesting sex-specific change in deregulation.

Since mirgenedb’s evolutionary analysis pinpoints which node in the phylogenetic tree a miRNA locus originates, we leveraged this information to date each miRNA locus. The distribution of all the miRbase annotations shows three peaks: at ∼10 million years, 150-200 million years and 500-600 million years [Figure 5h, Supplementary Table 8]. These correspond roughly to the speciation between macaque and mouse, the mid-Jurassic adaptive radiation in mammals^42^ and the Cambrian explosion^43^. Interestingly, the age deregulated miRNAs are more concentrated around 150-200 million and 500-600 million years [Figure 5i]. While this is the case for both males and females, the direction of regulation can differ. For example, for heart and prefrontal cortex, the direction of deregulation appears to be flipped, despite the magnitude of miRNA change being similar (between 25,000 and-25,000 rpmm) [Figure 5j].

With the ancient miRNAs being highly represented (peaks of 1500 for male and female while more recent miRNAs having peaks of ∼250 and 1000 respectively in Figure 5i) in terms of age deregulation, we wanted to see how many miRNAs within each family of miRNAs agreed between macaque and mouse. Even with comparable number of aging trajectories diminished because of tissue harmonization and homology constraints, the top families from Figure 2a are well represented. For example, MIR-154 and LET-7 still occupy the top 2 positions in terms of number of miRNAs deregulated. However, they differ in terms of how well they agree with the aging deregulation trajectories. Families such as MIR-154, MIR-17, MIR-92, MIR-15, MIR-133 appear to be skewed towards conservation of age deregulation between species, while LET-7, MIR-29 are either evenly split or skewed towards un-conserved directionality [Figure 5k].

We can highlight this mixed conservation of deregulation with the expression of the MIR-29 family in Figures 5l and 5m. The dominant version of the miRNA starts to differ as well as direction. E.g. In macaque monkey caudate putamen, mml-miR-29b-3p is the dominant form but is getting downregulated in females (changing by-9486 rpmm) while being upregulated in males (changing 12,320 rpmm). Meanwhile in mouse caudate putamen, mmu-miR-29a-3p is the dominant form and is being downregulated in females by-3022 rpmm and without detectable change in male mice.

## Discussion

Our dataset represents an unprecedented breadth of tissue, age and gender sampling for *Macaque mulatta* and *Macaque fascicularis* in comparison to previous studies focusing on a small number of tissues or blood^30,37^. Our detection of at least 400 miRNAs stably expressed across all tissues lines up with observations from humans and mice^44^. We are also able to successfully split the regions in our UMAP embeddings and pairwise clustering, suggesting successful tissue extraction and library preparation. However, we were tentative in interpreting the results from gallbladder due to high ratio of unique miRNA to aligned reads ratio.

While the monkeys were reared in a relatively controlled environment in comparison to being in the wild, there exist uncontrollable sources of variance. Individual variation is notable, especially in the non-brain organ samples. Among all the non-coding RNA types that are quantifiable using small RNA sequencing, the full length of mature miRNA molecules is captured by SE50 sequencing. MiRNA is also the best studied regulatory non-coding RNA class. Therefore, we restricted our analysis for this paper to balance the breadth of our dataset with depth.

Our findings agree with previous reports that miRNA deregulation with age does not overwhelm tissue-specific signatures. However, given the subtle change in expression over the ∼30 year lifespan of macaques, we were concerned about picking up technical noise as biological signal. With *M. musculus*, similar cohort size for each aging timepoints is possible, but this is not straightforward for larger long-lived organisms like macaques. Our sparse and continuous distribution of individuals reflects this. Therefore, we prioritized miRNAs that changed over a threshold [Supplementary Figure 1g] to maximize the probability of finding biological signals. Furthermore, we focus on families of miRNA that have been previously known to be functional, and used co-regulation based on genomic context to further filter out potentially spurious changes.

The lack of miRNA expression difference between *M. mulatta* and *M. fascicularis* is notable, since a difference is found even in different strains of mice. The two species of macaques have a DNA sequence similarity of 99.21% (approximately 22.7 Mbp difference out of 2.8 Gbp)^45^. Compared to ∼32.4 M difference in 13 strains of mice (1.2% of 2.7 Gbp)^46^. This suggests the genomic difference between the macaque species has not impacted the regulatory factors governing miRNAs, while between-mouse strain differences has. However, a deeper look at this is beyond the scope of this work. The similarity and limited age range of *M. fascicularis* led us to combine them together with *M. mulatta*.

The most frequently deregulated miRNA families in our dataset are known participants in aging. For example, miR-17-92 is one of the most frequently occurring families in pathogenesis with transgenic overexpression of miR-17 increased the lifespan of mice by ∼1.2 fold^27^. Downregulation of miR-17 in microglia (but not globally) of the Alzheimer’s model rescued spatial memory loss and anxious behavior associated^47^. The upstream transcription factor Lin28 of Let-7 can be upregulated to create optic nerve and spinal cord regeneration^48^, and in skin regeneration after thermal injury^49^. The MIR-181 family another example, with inhibition neuroprotective in a murine model of Parkinson’s^50^. What is striking is that deregulation of miR-181a in hippocampus is matching with macaques and *M. musculus* (increasing) but mismatching in other tissues. This highlights brain-region/matter specific control of miRNA and shows the pleiotropic nature of miRNA function in different tissues. A remarkable finding is that high fractions of the top 15 frequently deregulated miRNA families are also found in serum and plasma fractions in blood^38,51,52^. MiRNAs are generally abundant in blood cell types and extracellular vesicles^53^. Indeed, extracellular miRNA have been correlated with aging^54^ and could be a whole organism-wide signaling mechanism^55^. In our case, the only families with sizeable non-circulating members are MIR-15, MIR-29 and miRNAs without families.

Overall, there is a skew towards downregulation for miRNA aging trajectories. The proportion of miRNAs that are changing but don’t have any family suggests that a sizeable fraction of the change is likely noisy, with minimal impact on aging. The over-representation of miRNA families suggested a coordinated response. Notably, MIR-154 family is changing the most. It is genomically clustered inside the maternally imprinted *Dlk-Dio3* region^56^, one of the largest such clusters with 33 precursors inside a 0.8 Mbp region^57^. Interestingly, there are larger miRNA clusters (e.g. MIR-430 with 44 precursors known as the C19MC cluster^58^) that are not heavily deregulated, despite also being imprinted^59^. An estimated 61% and 66.2% of miRNAs are processed from host-genes in humans and mouse^60,61^, often within genomic clusters^10^, thus under control of surrounding regulatory elements / epigenetic states. The dramatic sex-tissue specific miRNA identity and direction of deregulation with age we observe in our data ties into this. The concordance of deregulation direction within miRNA clusters suggests that they are partially coordinated based on their upstream genomic and surrounding epigenomic context in a tissue-sex specific manner.

We sought to verify cross-species translatability of our macaque findings with our previous findings from *M. musculus*. While the mechanism of miRNA-Ago targeting is widely conserved, only ∼18 % of unique miRNAs are mappable by homology and synteny. Among homologous miRNAs, less than 50% have matching tissue of max expression. This diminishing conservation of miRNA by levels of organization (tissue, sex, miRNA cluster, individual miRNA, direction of deregulation) is also seen with aging. Differing concordance of within-cluster deregulation between brain regions and non-brain regions is conserved. So is deregulation direction flipping between the sexes, based on the tissue, with heart and full brain being notable examples [Supplementary Figure 3b]. Additionally, most age deregulated miRNAs are evolutionarily ancient. These together suggest that age deregulation is substantially controlled by upstream regulation rather than epigenetic drift. Based on this result, we might expect similar results in *H. sapiens* at the miRNA family level, but individual miRNAs will likely not translate well. The known difficulty in translating research from model organisms to human aligns with these findings ^62^.

While we have pointed out coordinated and noisy deregulation. Our most prominently deregulated MIR-154 family, inside the imprinted *Dlk-Dio3 cluster*^63,64^, is known to be associated with epigenetic clocks in *H. sapiens,* thus likely sits at the intersection point of both mechanisms. So far it has been studied in the context of aging. E.g. MEG3/Gtl2 (host gene for MIR-154 members) shows increased expression in hADSC prolonged passages^65^. Deregulation of members MIR-127 (chr7|+|14 in our dataset), another resident of *Dlk-Dio3* has also been linked to impaired neuronal survival^66^, appearing 34 times in multiple organs-tissue combinations in our dataset. In humans, it is also a potential biomarker of Frontotemporal dementia^67^ and found in circulating blood. MEG3-MIRG, the host gene for the miRNAs, have been associated with deregulation in aging in *M. musculus* heart^68^. MIR-430 (C19MC) is an even larger imprinted cluster of miRNA^69^, underlining *Dlk1-Dio3* as a hotspot unrelated to its size.

The prominence of this cluster’s deregulation in Brain regions and heart might be linked to the intensity of their metabolic activity (e.g. oxidative stress) interfering with the imprinting of these regions. However, widespread deregulation of this cluster throughout all brain matter types does not fully support the oxidative hypothesis; White matter has prominent deregulation, despite not being metabolically very active. Interestingly, inside of *Dlk1-Dio3*, the MEG3 perturbation has been implicated hippocampal apoptosis^70^, necroptosis in an Alzheimer’s disease model^71^, downregulation in Parkinson’s disease^72^, muscle regeneration ability^7^ and atherosclerosis^73^, with the direction of expression important for phenotypic outcome. However, the above interventions have excluded the MIR-154 cluster. While these miRNAs have been associated with pathogenesis^74,75^, studies are limited to muscle and blood^76^. The conservation of synteny with MEG3 in *H. sapiens, M. musculus and M. mulatta*^77^ suggests this cluster of miRNA might be a potential lever for regeneration, as has been demonstrated in skeletal muscle atrophy reversal not^78,79^. E.g. perturbation of miR-379-5p or miR-411-5p in a direction concordant with MEG3 perturbation from above studies in neural cell lines such LUHMES or Primary ventral mesencephalic and then testing for sensitivity to 6-OHDA stress would be a Parkinson’s relevant check.

Our study, of course, is not without limitations. A key challenge is the paucity of samples. Despite being one of the largest macaque aging cohorts, difficulty in having a cohort with multiple replicates for each age range meant we could not quantify within-age variance. Consequently, we prioritize the largest changes in miRNA expression using a data-informed inflection point cutoff threshold to filter out likely technical variation. This also influenced our ability to account for the impact of sex-specific hormones on life events. The modulation of sex differences through miRNAs is well known, e.g. sex-specific recovery after focal ischemia^80^, and regulation of QT interval in the heart^81^ along with sex-specific hormonal impact on aging^26,82^. While we captured strong sex-specific differences, the influence of individual life events such as menopause or puberty could not not be quantified with the current study design. Additionally, miRNAs-target interactions in rhesus macaques are still understudied. While evolutionary conservation of target sites is well studied, our lists of mRNA targets based on TargetScan likely contain false positives. Additionally, while we go into unprecedented depth in splitting up different brain regions, bulk RNA sequencing is unable to capture the cell-type specific richness within the regions or within other non-brain organs. Using emerging single-cell and spatial level non-coding RNA sequencing approach would represent an important next step, as would creating a validated miRNA-mRNA target set for macaques.

In conclusion, however, our dataset represents one of the most extensive rhesus monkey atlases from matching individuals. It adds an additional datapoint in the map between aging miRNA trajectories in mouse and humans, highlighting the largest clusters that are concordantly expressed and evolutionarily conserved between *M. mulatta* and *M. musculus*.

## Methods

### Macaque monkey rearing conditions

Rhesus macaques (*Macaca mulatta*) and cynomolgus macaques (*Macaca fascicularis*) included in this study were housed at the Biomedical Primate Research Centre (BPRC, Rijswijk, The Netherlands), a non-profit research institute maintaining a self-sustaining breeding colony of nonhuman primates established in the 1970s. Animals originate from captive-bred lines, with periodic introduction of new breeding lines to maintain genetic diversity and outbred status.

All housing and animal care procedures were conducted in accordance with European Directive 2010/63/EU and the Guide for the Care and Use of Laboratory Animals (NIH), and the facility is accredited by AAALAC International. Animal welfare and tissue collection procedures were reviewed internally by the institutional animal welfare body. No experimental interventions beyond routine veterinary care were performed on the animals included in this study.

Animals were socially housed in stable family groups of approximately 20–30 individuals, consisting of multiple matrilines and one adult male, reflecting natural social structures. Environmental enrichment was provided continuously. Animals were fed a commercially available primate diet (SSniff, Germany) supplemented with fruits, vegetables, and grains, with water available ad libitum.

Animals were monitored daily by trained caretakers, with health assessments including behavioral observations, injury checks, and evaluation of general condition. In addition, animals underwent routine veterinary examinations, including yearly blood sampling for haematological and biochemical analyses, as part of the health monitoring program.

Tissue collection for this study was performed within the framework of the BPRC Nonhuman Primate Brain Bank (NHPBB)^83^. Postmortem procedures were carried out under standardized conditions with a short postmortem interval (∼1 hour), ensuring high-quality preservation of RNA and tissue integrity for downstream molecular analyses.

### RNA library Preparation and sequencing

Total RNA including miRNAs was isolated from tissue sections using miRNeasy Mini Kit (Qiagen, Hilden, Germany) according to the manufacturer’s recommendations. As previously described1, isolated RNA was purified by ethanol precipitation followed by additional clean-up of the RNA samples using the RNeasy Clean Up Kit (Qiagen, Hilden, Germany). RNA concentration was determined using the NanoDrop™ 2000 spectrophotometer (Thermo Fisher Scientific, Waltham, MA, USA) and RNA integrity of all samples was assessed using the Agilent RNA 6000 Nano Kit for Agilent 2100 Bioanalyzer (Agilent Technologies, Santa Clara, CA, USA). smallRNA libraries were prepared on the high-throughput MGISP-960 sample preparation system using the MGIEasy smallRNA Library Prep Kit (MGI Tech Co. Ltd., Shenzhen, Guangdong, China). In brief, 100 ng of total RNA was used as input. After adapter ligation, reverse transcription. PCR amplification and size selection, concentrations of amplified PCR products were measured using Qubit™ 1x dsDNA HS Assay Kit (Thermo Fisher Scientific, Waltham, MA, USA). For each library, 20 to 21 different barcoded samples were pooled and circularized to generate single-stranded DNA libraries (ssDNA), and the concentration of the ssDNA libraries was determined using Qubit™ ssDNA Assay Kit (Thermo Fisher Scientific, Waltham, MA, USA). Subsequently, ssDNA libraries were used to generate DNA nanoballs (DNB) by rolling circle amplification. Libraries were sequenced using the DNBSEQ-G400RS High-Throughput Sequencing Reagents Set SE50sR on the respective instrument.

### Non-coding RNA quantification

Fastq files were quantified using the mirmaster 2.0 pipeline^84^ as follows: the adapter “AGTCGGAGGCCAAGCGGTCTTAGG” was used for trimming with a minimum overlap requirement of 10 nucleotides with a maximum edit distance to adapter of 1 using a sliding window size of 4 nucleotides. A quality threshold of 20 also had to be met inside the sliding window. If the tail or head of a read had 3 ambiguous reads, they were trimmed. No ambiguous nucleotides were allowed in the trimmed reads. Only reads at least 17 nucleotides long were kept. Alignment was performed using bowtie ^85^ which allows for a single mismatch using the following options: bowtie-m 100 –best –strata –fullrefs. For *M. mulatta and M. fascicularis* mmul_8.0.1 with the NCBI assembly ID GCA_000772875.3 was used as alignment. For *M. musculus* GRCm38 with NCBI assembly ID GCA_000001635.2 was used. Mirmaster 2.0 relies on the miRbase annotation format, and here miRbase v22^86^ was used for miRNA quantification. Since many of recent miRbase miRNA additions have a higher probability of false positives we additionally used mirgenedb 3.0^10,86^ to prioritize annotated miRNAs with a higher probability of functionality. Quantification of other non-coding RNA were performed using GtRNAdb v18.1^87^, Ensembl ncRNA v100 ^88^ and NONCODE v5^89^.

For *M. mulatta* Novel miRNAs were quantified based on miRMaster 2.0 output and subject to further filtration by stable expression with at least 5 raw counts in 50% of samples in a tissue. Furthermore we looked at the RNAfold^90^ generated structure of the precursors to check for hairpin structure, 5’ overhang, along with pileup plots generated from the aligned BAM files to check for reduced 5’ heterogeneity^91^.

For all non-coding RNA, we filtered for stably expressed miRNAs with 5 raw counts in at least 50% samples in at least one non-brain organ / brain region. Reads per million mapped reads normalization (rpmm) was used across all non-coding RNA to account for variances in sequencing depth and maintain comparability.

### Circulating miRNA determination

Lists of macaque circulating miRNA were obtained from the supplementary data of the following papers^38,51,52^. Selection criteria of miRNAs from the paper is dependent on the paper. Since Schneider et. al. ^51^ looks at plasma miRNAs in fasting rhesus monkey, we keep all miRNAs reported to be detected. Chen et. al.^52^ report plasma miRNAs before and after ischemic stroke is reported and we keep the miRNAs reported as non-zero mean values I the after-stroke group. Russ et. al. looks at serum miRNA before and after radiation exposure is reported. Since heavy radiation exposure is not common with extended age, only miRNAs in 50% of the samples in the 96 hours pre-radiation group were included.

### Tissue specificity index calculation

For *M. mulatta* group, tissue specificity index was calculated using the following equation:

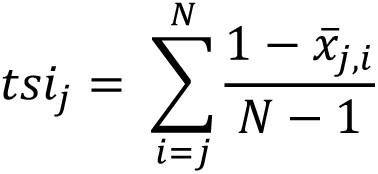

Where j is an ncRNA, i is an organ, N is the total number of organs in the analysis and *x_j,i_* is the mean log2 rpmm scaled expression for the ncRNA j for the organ i. Since brain is the only organ with within-organ values, the labels of different brain regions were collapsed into a single “brain” label to account for within brain-region similarity artificially decreasing the TSI values.

Tissue specificity index values of *M. musculus* were obtained from the miRNATissueatlas 2025^21^, where TSI values are calculated similarly.

### Aging trajectory calculation

Due to the dearth of sampling density across the approximately 30 year lifespan of rhesus macaque, we opted a multi-pronged approach to finding deregulated non-coding RNAs. We calculated the spearman rank correlation, a non-parametric test of correlation, and filtered by a correlation coefficient of 0.3.

Since Spearman rank is the covariance of two variables (here age and rpmm expression) normalized by their standard deviations, the absolute value is not necessarily the effect size of age on rpmm. Therefore, we fit an ordinary least square (OLS) equation between age and miRNA rpmm, using the change over the lifespan as a proxy for effect size. We plotted the distribution of the OLS-predicted expression changes and based on expression change value where the frequency went through the greatest change, picked thresholds of +10 and-10 rpmm to filter for the changes with the largest effect size.

### Fitting linear mixed models for variance estimation

To quantify the variance contributed to each non-coding RNA’s expression, we fit two linear mixed models. Two distinct models were used as the individuals were distinct between the cohorts were distinct.

Non-brain organs:

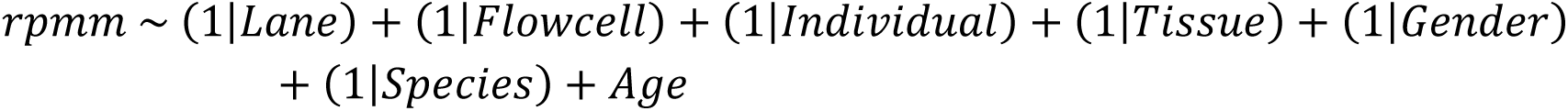

Brain regions:

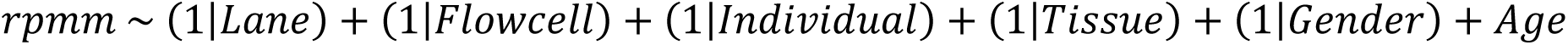

### miRNA cluster concordance calculation

The miRNA genomic coordinates from miRbase were used to assign genomic clusters if miRNAs were within a 50 kilobase window of each other and arranged in the same direction. The nomenclature is “chr|+|integer” where chr represents the chromosome on which the cluster is found, positive or negative sign indicates the read direction and the integer is an arbitrary number assigned to each cluster.

Concordance distances of miRNAs were calculated based on the OLS expression change across lifespan for pairs of miRNA according to the following equation:

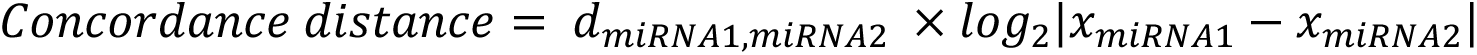

Where 𝑑_𝑚𝑖𝑅𝑁𝐴1,𝑚𝑖𝑅𝑁𝐴2_ is +1 if the OLS direction change is matching between miRNA1 and miRNA2, and-1 if OLS direction change is mismatching. 𝑥_𝑚𝑖𝑅𝑁𝐴𝑛_ refers to the OLS direction change for miRNA n calculated as in “Aging Trajectory Calculation” subsection.

### miRNA pathway analysis calculations

For *M. mulatta*, currently there is no large set of experimentally validated miRNA-target interactions available. Therefore, we had to rely on predicted seed-target hits that are highly conserved in the Targetscan database^15^. Furthermore, we filtered using a 75% context score threshold to reduce the likelihood of false positives.

Using the list of targets, we used clusterprofiler^92^ to perform overrepresentation analysis using the enrichKEGG function^93^. The background universe was set as the overlap of mapped targets from Targetscan and Entrez gene names. The parameters used were minGSSize of 3, maxGSSize of 500, with a Benjamini-Hochberg pAdjustMethod. An adjusted pvalue cutoff of 0.05 was applied. We report the unaggregated pathways left with this cutoff along with pathways aggregated by subcategory, to reduce the number of pathways in our graph and provide a more coarse overview. For the aggregation, the number of genes on each pathway inside each subcategory were summed together, while the negative log10 of pvalues are logged.

### miRNA homology and evolutionary age calculation

To assign the years when a miRNA precursor locus originated, information from mirgenedb 3.0 and TimeTree 5^94^ was used. For the list of taxonomic names list in the “Node of origin (locus)” column of mirgenedb entries, we assigned a taxonomic id (taxid) by using the Python script accessing the eutils API from NCBI^95^. The taxid list was used with the TimeTree 5 API (accessed August 25 2025) to access the corresponding branch length of the node. The branch node corresponds to Millions of Years Ago (MYA). Since we also map mirgenedb names to miRbase names which contain miRNAs that have not been assigned to homologues, we make the assumption that these miRNA loci are species-specific are as old as the species itself with values of 6.19 MYA for *M. mulatta* and 6.77 MYA for *M. musculus*.

### Software

Snakemake^96^ 8.25.4 and Conda 23.1.0 was used to create the workflow pipeline and manage packages. Scripts were written using Python 3.14.2 and R 4.4 The following Python packages were used: Numpy 2.3.5, Pandas 2.3.3., Scikit-learn 1.8.0, Scipy 1.16.3, Statsmodels 0.14.6, Pyarrow 21.0.0, Umap-learn 0.5.11, Scanpy 1.12, gff3 1.0.1, biopython 1.85, beautifulsoup4, html5lib, requests, lxml. The following R packages were used: ggplot2 4.0.1, complexheatmap 2.26.1, dplyr 1.2.0, svglite 2.2.2, dendextend 1.19.1, ggplotify 0.1.3, aplot 0.2.9, tidyr 1.3.1, ggtree 4.0.4, ggrepel 0.9.6, ggh4x 0.3.1, ggdendro 0.2.0, cowplot, upsetr 1.2.0, ggupset 0.4.1, tidyverse 2.0.0. Literature research, and code syntax lookup were streamlined using Claude Sonnet 4.6.

## Data and Code availability

Raw and processed data are available under the GEO accession GSE328747. Code will be made available upon request.

## Author Contribution

S.R.1. performed conceptualization, formal analysis, visualization and writing-original draft.

E.N. performed sample collection and project administration. N.L., L.G. performed library preparation and project administration. T.T. performed library preparation. U.F., S.R.2., M.H. performed validation experiments. J.M., M.F. performed formal analysis and visualization. M.A.E. performed conceptualization, supervision, writing-review and editing.

J.M. performed conceptualization, project administration, supervision, writing-review and editing. A.K. performed conceptualization, funding acquisition, project administration, supervision, writing-review and editing.

## Funding and Licensing Statement

This project was funded by Michael J Fox Foundation grants MJFF-028018, MJFF-021418, Deutsche Forschungsgemeinschaft (DFG) grant KE 1894/20-1. Figure 1a was generated with Biorender, license No. VI29LLK1BN.

## Declaration of interest

The Authors declare no competing interests.

## Declaration of generative AI and AI-assisted technologies in the writing process

During the preparation of this work the author(s) used Claude Sonnet 4.6 to streamline literature searching and code syntax lookup. After using this service, the author(s) reviewed and edited the content and needed and take(s) full responsibility for the content of the published article.

## Acknowledgements

All caretakers and veterinary staff are sincerely acknowledged for their assistance during the annual health exams and for their exceptional care for the animals. The start-up of the non-human primate brain bank is in part supported by a Longevity Impetus Grant from Norn Group, Hevolution Foundation and Rosenkranz Foundation.

---

**Supplementary Table 1:** Per sample QC metrics for small RNA sequencing runs

**Supplementary Table 2:** Aging trajectory of macaque non-brain organs and brain organs, split by sex.

**Supplementary Table 3:** Tissue specificity index of macaque organs for males, females and combined. Brain regions have been collapsed to a single organ to prevent artificial deflation of TSI values.

**Supplementary Table 4:** Macaque miRNA deregulation overlap truth table for non-brain organs males.

**Supplementary Table 5:** Macaque miRNA deregulation overlap truth table for non-brain organs females.

**Supplementary Table 6:** Macaque miRNA-target pathway analysis result per brain-region and non-brain organ

**Supplementary Table 7:** Pairwise distance between within cluster miRNAs and between cluster miRNAs for non-brain organs and brain regions within each sex for macaque. MiRNA clusters assigned based on a 50 Kb window of the genome and genomic direction.

**Supplementary Table 8:** MiRNA homology assignment based on mirgenedb. Evolutionary age calculated from TimeTree is also included.

**Supplementary Table 9:** Tissue-specific index for miRNA homologous between *M. mulatta* and *M. musculus*

**Supplementary Table 10:** Pairwise distance between within cluster miRNAs and between cluster miRNAs for non-brain organs and brain regions within each sex for *M. musculus*. MiRNA clusters assigned based on a 50 Kb window of the genome and genomic direction.

**Supplementary Figure 1:**
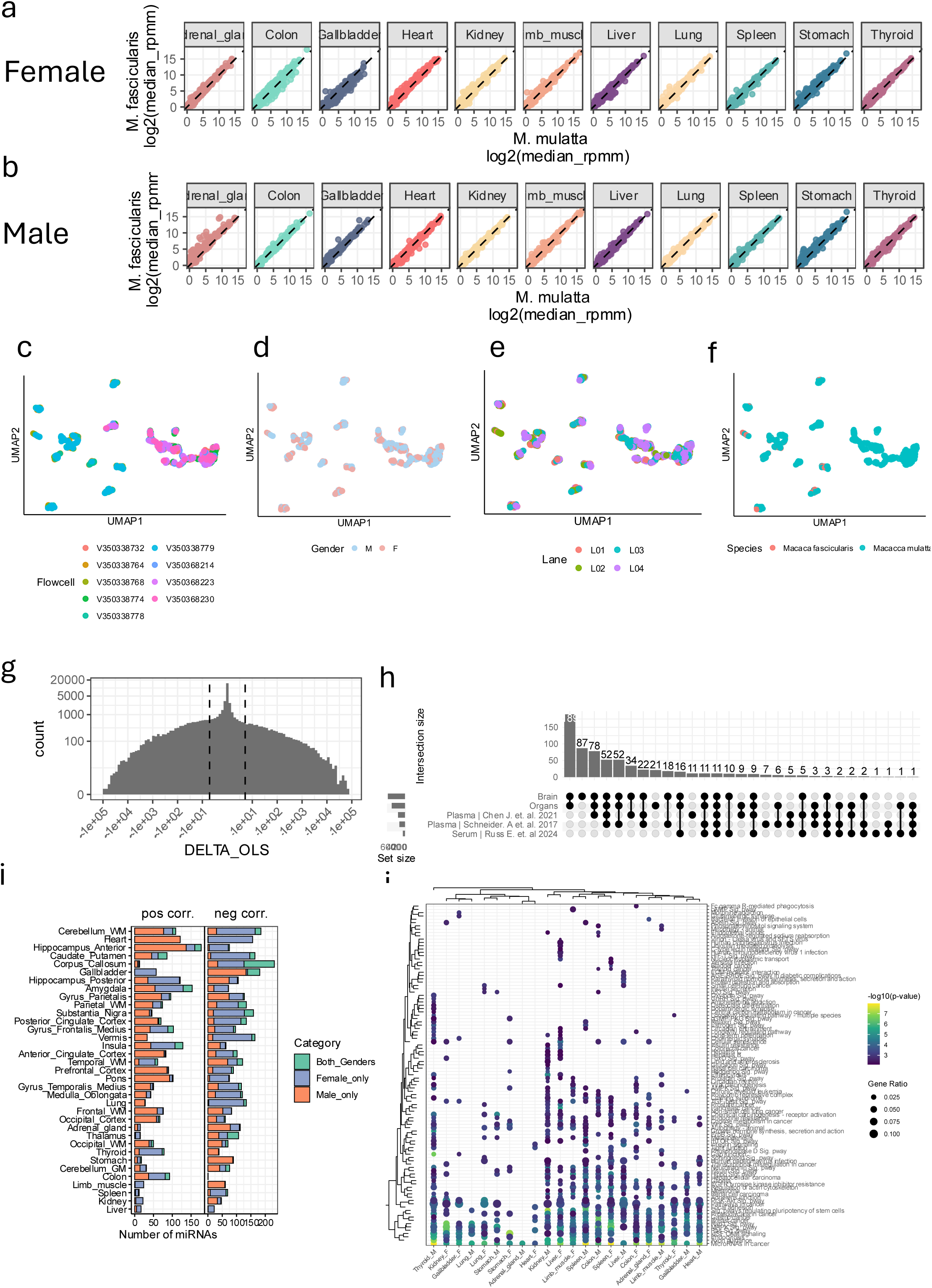
a) Comparison of median miRNA expression between *M. mulatta* and *M. fascicularis* females. b) Comparison of median miRNA expression between *M. mulatta* and *M. fascicularis* males. c) UMAP embedding colored by Flowcell, d) Gender e) Lane on Flowcell f) Rhesus Macaque species. g) Distribution of the frequency of expression change as calculated from OLS fit for each miRNA for each tissue for each gender. Count is log transformed. Threshold used as cutoff for likely-technical noise is represented using dashed lines. h) Overlap between the miRNAs detected in brain, non-brain organs and blood datasets from literature. i) Frequency of miRNAs for brain regions and non-brain organs, colored based on if they’re found only in male, only in female or shared between the genders. Frequencies are further split based on if the direction of change is positive(pos corr.) or negative (negative corr) j) Pathway overrepresented by targets of deregulated miRNA in each tissue in each gender for only non-brain organs.

**Supplementary Figure 2:**
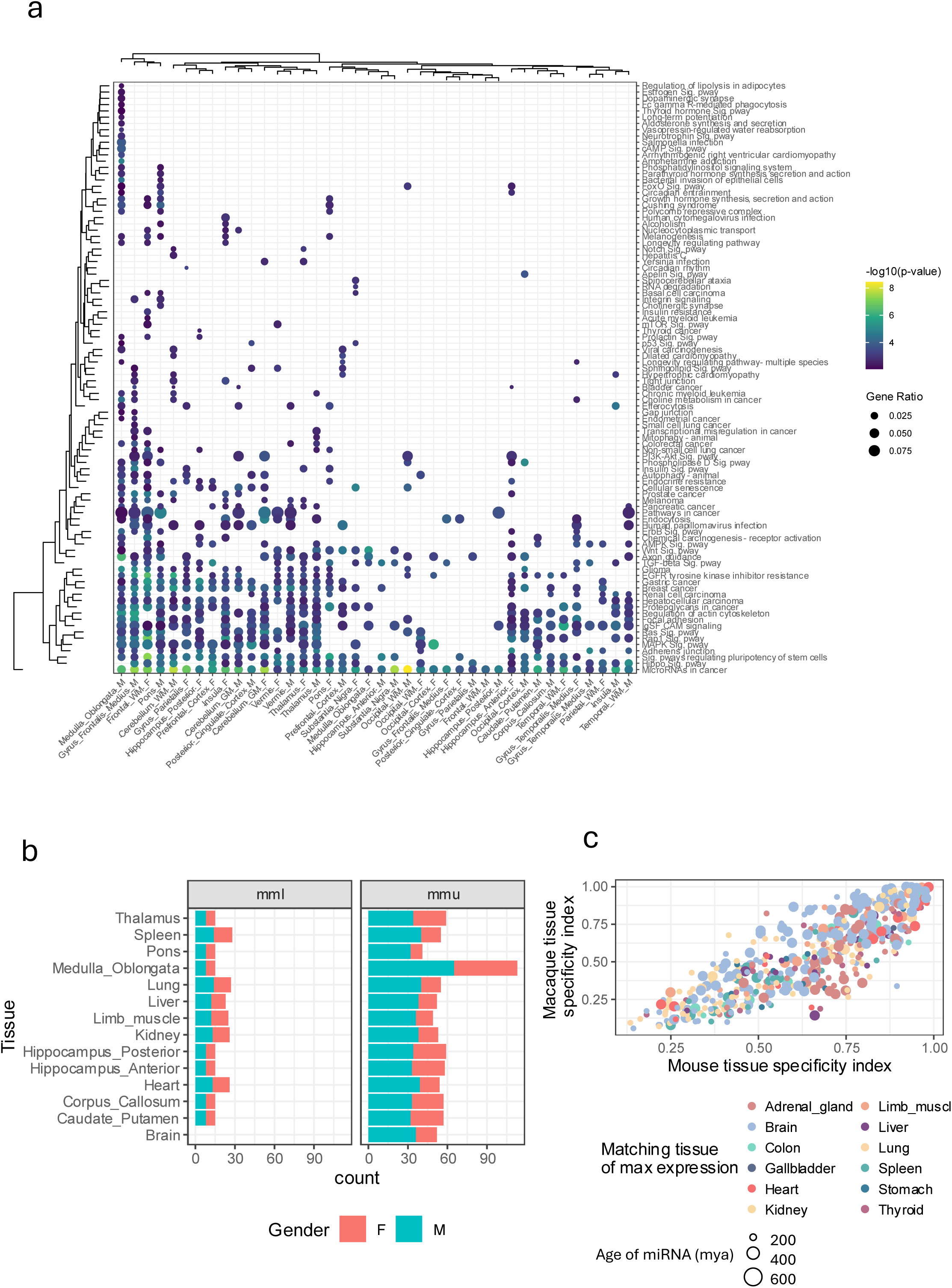
a) Pathway overrepresented by targets of deregulated miRNA in each tissue in each gender for only brain regions. b) Number of samples for each gender of each species based on this macaque dataset and previous Tabula Muris senis non-brain organs and brain regions dataset. Only harmonized tissues compared, apart from Brain c) Homologous miRNA comparison between macaque and mouse, colored by tissue of max expression in macaque monkey. Size of the dot represents the evolutionary age.

**Supplementary Figure 3:**
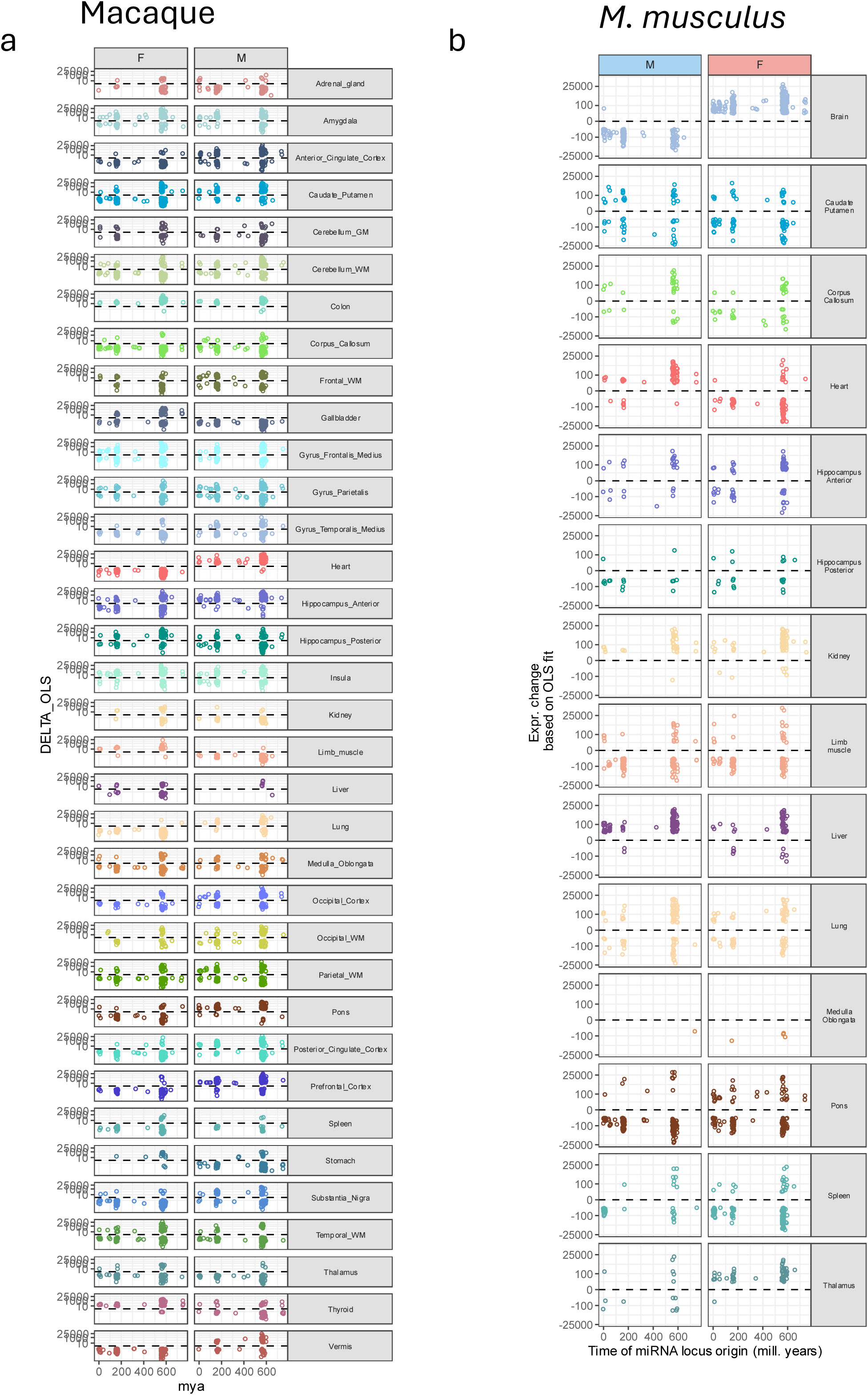
a) OLS derived expression change for each miRNAs trajectory for each tissue in each gender against evolutionary age for macaque monkeys. b) OLS derived expression change for each miRNAs trajectory for each tissue in each gender against evolutionary age for *M. musculus*.

## Notes

### Competing Interest Statement

The authors have declared no competing interest.

### Summary of Updates

First line of abstract has been adapted to highlight the importance of studying the molecular biology of aging. Certain connecting words have been modified in the rest of the abstract to stay within a word limit of 150. Orchid ID has been corrected.

